# Protocols for rational design of protein solubility and aggregation properties using Aggrescan3D standalone

**DOI:** 10.1101/2020.09.09.276915

**Authors:** Aleksander Kuriata, Aleksandra E. Badaczewska-Dawid, Jordi Pujols, Salvador Ventura, Sebastian Kmiecik

## Abstract

Protein aggregation is a major hurdle in the development and manufacturing of protein-based therapeutics. Development of aggregation-resistant and stable protein variants can be guided by rational redesign using computational tools. Here, we describe the architecture and functionalities of the Aggrescan3D (A3D) standalone package for the rational design of protein solubility and aggregation properties based on three-dimensional protein structures. We present the case studies of the three therapeutic proteins, including antibodies, exploring the practical use of the A3D standalone tool. The case studies demonstrate that protein solubility can be easily improved by the A3D prediction of non-destabilizing amino acid mutations at the protein surfaces.

## 1. Introduction

Protein-based therapeutics, such as antibodies, receptor decoys and replacement enzymes have already transformed the drug discovery field and have the potential to yield new therapies. Perhaps the most difficult challenge facing development and manufacture of protein therapeutics is protein aggregation^1^. The experimental identification of protein regions responsible for aggregation can be expensive and time-consuming. Therefore, there is an increasing need for computational tools that can support these efforts and aid in the design of soluble protein variants^1^.

In 2015, we introduced Aggrescan3D (A3D) web server for prediction of aggregation propensity in protein structures^2^. The method relies on experimentally-derived scale of aggregation propensity for each natural amino acid^3^, which has been successfully used in a sequence-based prediction Aggrescan method^4^. In addition, A3D uses structure information and allows for the identification of aggregation-prone residues on protein surface. Through introducing virtual mutations, A3D is an effective tool for the design of protein variants with increased solubility and testing of the impact of pathogenic mutations. A3D server enables also to take into account the dynamic flexibility of protein structures, which may influence aggregation propensity. This is possible in A3D Dynamic Mode that uses the CABS-flex method for the fast simulations of flexibility of globular proteins^5,6^. Since 2015, A3D has been successfully used in many studies aimed at enhancing protein solubility^7–10^.

In 2019, following the users requests, we introduced the updated 2.0 version of the A3D web server^11^ and A3D standalone application package^12^. The A3D 2.0 web server offers several new functionalities like simultaneous prediction of changes in protein solubility and stability upon mutation, automated mutations tool that identifies high scoring residues and suggests protein variants with optimized solubility, dynamic mode calculations for large and multimeric proteins, a REST-ful service to incorporate A3D calculations in automatic pipelines, and a new, enhanced web server interface. The A3D standalone application is a fully functional implementation of the A3D 2.0 web server that is intended to work locally and introduces command line utilities. Therefore, A3D standalone addresses the important aspects of data privacy and user control over every stage of the modeling process.

In this work, we briefly describe the A3D standalone functionalities (see section 2), provide some basic examples of its usage (see section 3) and demonstrate A3D applications for rational design of antibodies and other therapeutic proteins (see section 4). As demonstrated, the A3D standalone is a flexible tool that can be adapted to a variety of needs and easily incorporated by the users into their own pipelines.

## 2. A3D standalone package

### 2.1. Installation instructions and requirements

Local installation of A3D standalone requires Python 2.7. We recommend using Anaconda - a scientific Python distribution that comes with pre-installed packages and conda package manager which allows for an easier and less error prone installation. Installation of Anaconda is straightforward. For this purpose, download the installer for Python 2.7 from their web page and follow the instructions below (though A3D can be installed in many ways, more instructions are provided in the A3D online repository, see Note 1).

The installation of base A3D functionality can be achieved with one command (note: Windows users should use the Anaconda Prompt):

*$ conda install -c lcbio aggrescan3d*

To verify the installation run:

*$ aggrescan -i 2gb1 -w test_run -v 4*

A3D comes with a built-in server-like app that can be operated through the web browser (we recommend Google Chrome and suggest not to use Internet Explorer as it will not support full functionality of the app). To verify the app installation type:

*$ a3d_server*

On the first run you will be prompted to install packages needed for the server. Type ‘y’ to install them - this is mandatory for the app to run. Re-run the command - you will be prompted for the FoldX location. We recommend downloading and providing FoldX’s location at this point, although it can be set up later via the UI. If the installation succeeds, the server should start and be available under the address *localhost:5000* in your web browser. Please note that the A3D program must continue running for the web server to function.

A3D has two main dependencies that are needed for its basic functionalities: FoldX^13,14^ and CABS-flex packages^5^ (see the A3D pipeline in Figure 1). FoldX is used for the energetic minimization of input structures, creation of mutant structures and stability calculations. CABS-flex standalone package is required for the dynamic mode to generate a set of models reflecting protein flexibility. Both packages can be downloaded free of charge for academic users (for download instructions see Note 1).

**Figure 1.**
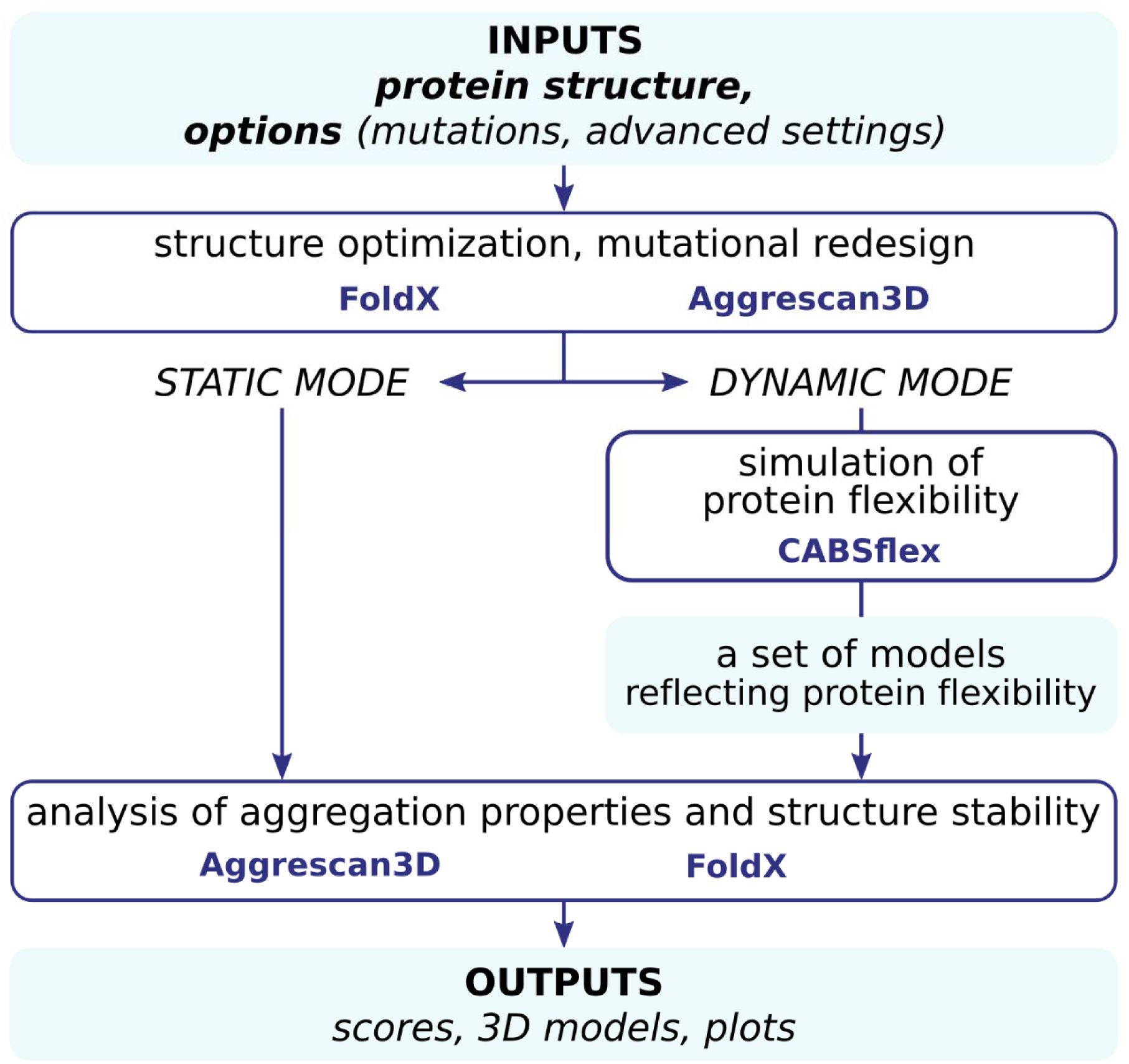
Aggrescan3D pipeline.

### 2.2. A3D input commands

The only required input of the A3D is a protein structure in PDB format. The Table 1 introduces basic input and set-up commands. Additional technical tips are provided in Note 2.

**Table 1.** A3D input and setup commands.

| *command* | *description* | *example* |
| --- | --- | --- |
| -i, --protein<br>PDB | Allows providing input protein structure as PDB code (fetched from *www.rcsb.org*) or a text file in pdb or pdb.gz format. | -i 2GB1<br>-i *filepath* |
| -C, --chain<br>CHAIN_ID | Limits the A3D analysis to the specific protein chain(s) provided as the valid chain identifiers. All other chains will be ignored. | -C A |
| -c, --config_file<br>CONFIG_FILE | Allows providing a configuration file that sets up the program's parameters. Note that commands from command line override the config file. | -c *filepath* |
| -D, --distance<br>VALUE | Sets up the distance value used in A3D score calculations (see section 3.1). *default*: 10 | -D 10 |
| -f, --foldx | Introduces a structure repair by FoldX program before A3D stage. The flag is also required when mutation options are used. | -f *filepath*<br>-f |
| -n, --naccess | Uses naccess instead of freeSASA for solvent-accessible surface area calculation. | -n |

### 2.3. Mutation commands

A3D allows testing the effect of mutations on the input structure. The mutations can be introduced (i) manually, where the user specifies a residue and a mutation type; or (ii) automatically on the most aggregation prone residues identified by A3D analysis. Mutants are generated using the FoldX program (*--foldx* option is required) and A3D analysis is performed on each mutant including the calculation of energetic effect of the mutation. The Table 2 introduces mutation commands.

**Table 2.** A3D mutation setup commands.

| *command* | *description* | *example* |
| --- | --- | --- |
| -m, --mutate<br>VALUE | Provides the mutation(s) code applied to the input protein. The syntax for the option:<br>**-m** <old_residue><new_residue><residue index><chain ID><br>See Note 2 for additional tips. | -m MW1A |
| -am, --<br>auto_mutation<br>OPTIONS:<br>N M [] | Predicts more soluble mutants (into Glu, Asp, Arg, Lys) for *N* automatically selected residues. The second argument, *M*, is the number of used cores. An optional argument is a list of residues for which mutations are excluded. The syntax for the option:<br>**-am** '<max_number_of_mutated_residues><br><number_of_cores_used><br>[list_of_excluded_residues as <residue ID><chain ID>]'<br>See Note 2 for additional tips. | -am '2 2 1A 2B 3C' |

### 2.4. Dynamic mode commands

In dynamic mode, A3D uses CABS-flex tool^5^ for fast simulations of protein flexibility. CABS-flex generates several protein models, which reflect protein flexibility^15,16^, and a full A3D analysis in static mode is performed on each of the model. A3D takes full advantage of the CABS-flex package and allows the user to tailor the simulation to their needs. Table 3 describes the dynamic mode commands.

**Table 3.** A3D dynamic mode commands.

| *command* | *description* | *example* |
| --- | --- | --- |
| -d, --dynamic | Uses the CABS-flex tool for the protein flexibility simulation and aggregation propensity analysis on generated models. | -d |
| --n_models<br>VALUE | Chooses the number of models that CABS-flex will generate for the analysis. | --n_models 8 |
| --cabs_dir<br>PATH | Specifies a path for CABS-flex program. This is mostly intended as a developer option. | --cabs_dir<br>/PATH/CABS |
| --cabs_config<br>PATH | Specifies a path for a CABS-flex configuration file that gives the user more control of the CABS-flex simulation. | --cabs_config<br>/PATH/CABS_config.ini |

### 2.5. Output commands

Output commands mostly concern logging and debugging as the calculations output remains much the same. We recommend using a high verbosity value to gain insights on how the program operates. Table 4 provides the output-related commands.

**Table 4.** A3D output commands.

| *command* | *description* | *example* |
| --- | --- | --- |
| -v, --verbose<br>VALUE | Decides how much information will be reported during the program run. Choose between: **0** - *Critical*, **1** - *Warning*, **2** - *Info* (default), **3** - *Log files*, **4** - *Debug*. See Note 2 for additional information. | -v 4 |
| -r, --remote | Disables the display of logging to the console and stores the logs in the working directory in the *Aggrescan.log* file instead. | -r |
| -M, --movie<br>FORMAT | If PyMol is installed a short movie of the protein is saved. The supported formats are 'webm' and 'mp4'. | -m webm |
| -w, --work_dir<br>PATH | Saves all the results and temporary files to the specified PATH (note: this will overwrite existing results if the same path is specified multiple times) | -w test_run |

## 3. Examples

### 3.1. Basic run in static mode

Once the installation is done, A3D can be used in two ways: the command line or the app. Please note that for Windows all the commands need to be run with Anaconda Prompt. Below, we describe how to run a simple job using both methods.

Using command line run (for the command syntax see Table 1 and 4):

*$ aggrescan* -*v 4 -w a3d_simple -i 2gb1*

This command keeps the job in an *a3d_simple* folder, where one can check all the output files provided by A3D:

- *A3D.csv* **-** text file with A3D score results consists of columns: (1) *Protein* - an internal name used by the program, (2) *Chain* - one-letter chain ID, (3) *Residue* - residue’s index, (4) *Residue_name* - one-letter code identifying the amino acid, (5) *Score* - Aggrescan3D score;
- score plot for each chain in PNG format, *chainID.png*;
- PDB files **-** *input.pdb* and *output.pdb* (the result of the simulation with b factor field replaced with A3D score);
- *config.ini* **-** configuration file that allows the user to re-run the simulation with the same settings or changing some options (see Note 2).

To run the same simulation in the app, run the server (*a3d_server*) and don’t close that terminal window. Go to *localhost:5000* in your browser and use the interface to set up the parameters of the run. Click ‘Run’ button to start the program. We provide the sample setup in the Figure 2. The example of using dynamic mode is provided in After the run ends, A3D results will be displayed in several tabs, providing the details of the prediction and several interactive tools to inspect A3D scores and visualize the protein structure. User’s experience when running the A3D app should be similar to our A3D 2.0 web server. For the analysis of the results:

**Figure 2.**
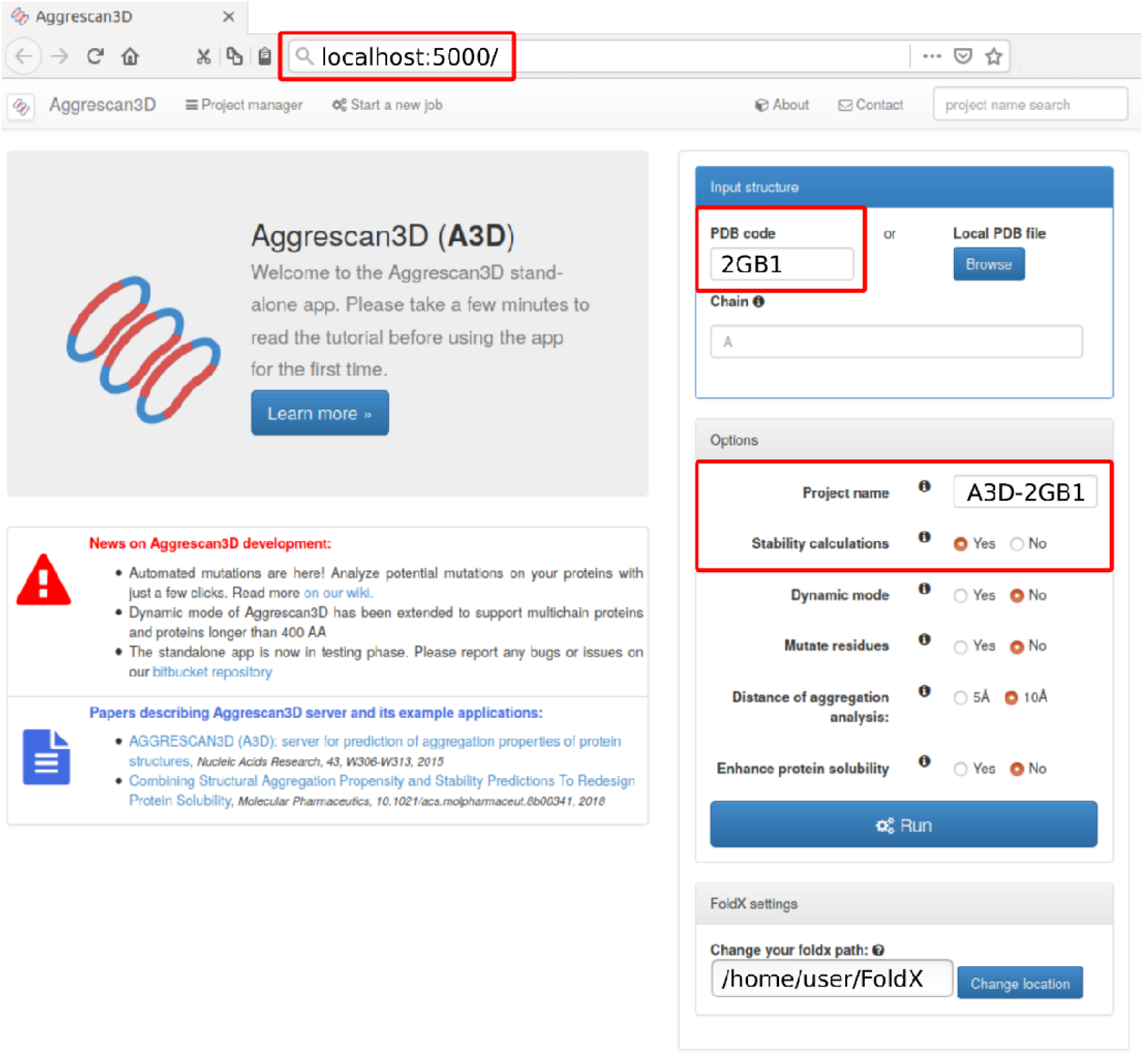
Web interface of the A3D standalone. The customization of the simulation run may be set within ‘Input structure’ and ‘Options’ panels marked in red.

- *Plot* tab presents the interactive per-chain plots of A3D scores for each residue;
- *Score* tab provides score statistics and a score overview for each residue in tabular format;
- *Structure* tab provides a visualization of the protein structure with coloring based on the A3D score. Residues can be labelled, and snapshots taken which are all available in the *Gallery* tab. The example results are presented in the Figure 3.

### 3.2. Automated mutations run

The A3D automated mutations feature automatically invokes 2 consecutive modeling steps. In the first, A3D analyses the input structure in the static mode and identifies the most aggregation prone residues (2 by default). In the second step, A3D creates mutants in which the most aggregation prone residues are mutated to each of the 4 charged amino acids: D, E, K and R (if 2 aggregation prone residues are identified in the first step then 2×4=8 mutants are created). It has been demonstrated that the presence of charged residues attenuates the aggregation propensity of hydrophobic stretches by provoking electrostatic repulsions, thus acting as gatekeepers of aggregation^21^. The automated mutations feature (-*am* option) creates a ranked list of point mutations, where both the solubilizing (AvgScore, AvgScoreDiff) and energetic (EnergyDiff, ΔG) effects are considered. More negative A3D average scores point to higher solubility predictions, while negative energetic effects suggest thermodynamically stabilizing mutations.

**Figure 3.**
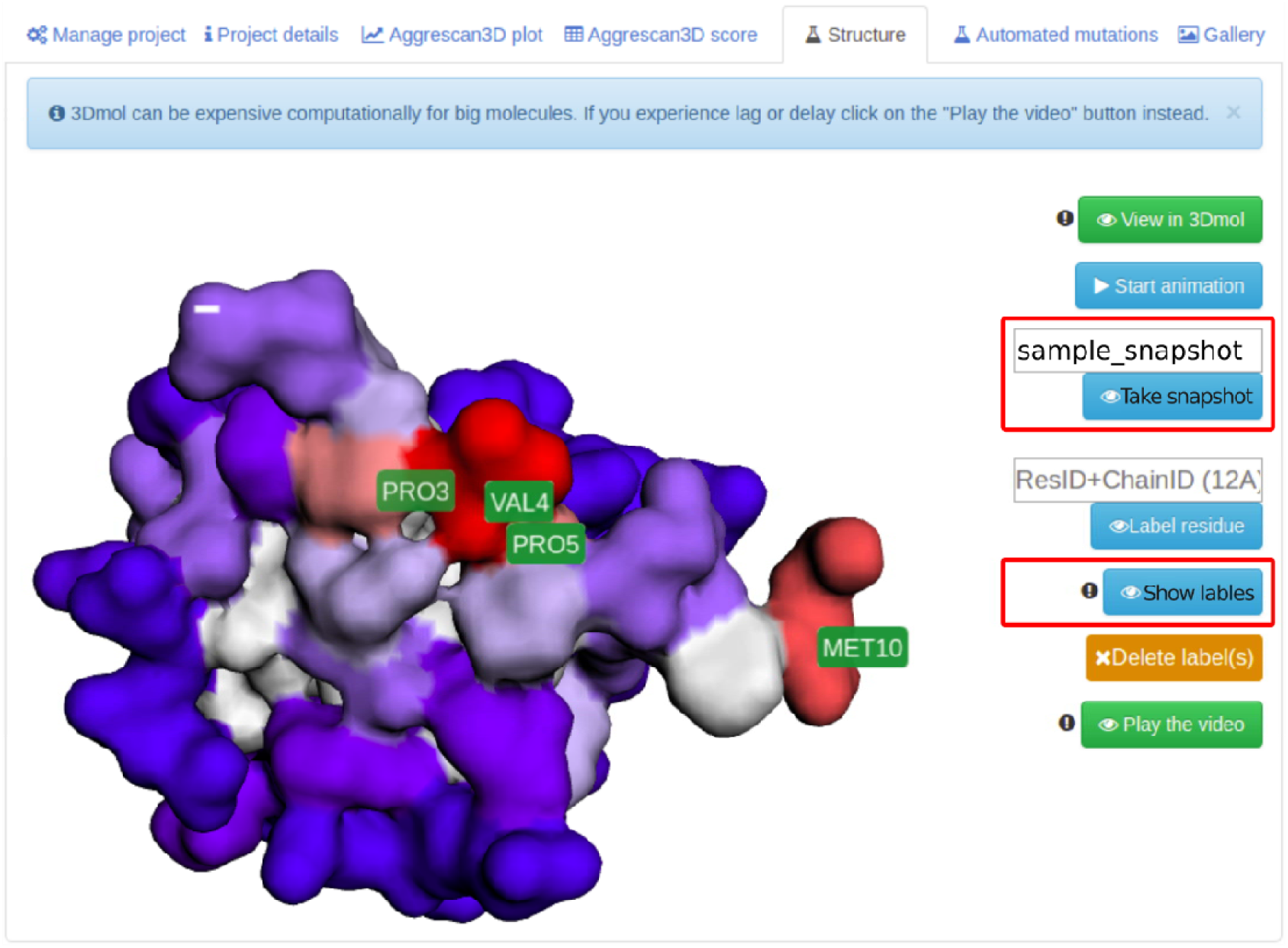
The example protein structure analyzed in the *Structure* tab of web interface of the A3D standalone. Use the *Show labels* button to select labels for aggregation prone residues. Click *Take snapshot* to save the current view in the *Gallery* tab. The protein structure can be freely rotated with holding left mouse button and zoomed in and out with the mouse wheel.

This feature requires providing a path to FoldX force field algorithm. Using command line run (the command syntax is described in Table 2):

*$ aggrescan -v 4 -w a3d_enhance -i 2gb1 -am -f /path/to/foldx*

This will generate several additional result files (the format of mutation codes is explained in Note 3):

- *Mutations_summary.csv* - CSV file containing the energetic and solubility effect for each created mutant;
- <mutation_code>.*csv* - CSV file with A3D scores of each residue for the specific mutant;
- <mutation_code>.*pdb* - PDB output file with the mutant’s structure;
- <short_code>.*png* - plot of A3D score for each mutant of the specific residue with the wild type as a baseline (PNG format);
- <Short code>.*svg* - plot of A3D score for each mutant of the specific residue with the wild type as a baseline (SVG format).

To run the same simulation in the app, select *Enhance protein solubility* and *Stability calculation* options in the app interface (see Figure 2, the interface enables also providing a path to FoldX force field). After clicking ‘Run’ button a new window will appear where certain amino-acids can be frozen i.e. excluded from the mutation procedure. To run the program click ‘Save changes and submit’. A3D models multiple mutants with solubilizing substitutions in the aggregation-prone regions of the initial structure. These structures are re-analyzed, and the predictions reported in the new tab *Automated mutations*. There, users can compare the impact on the aggregation propensity (A3D score) and protein stability (ΔG) for each of the mutants and visualize their 3D structures (see Figure 4).

**Figure 4.**
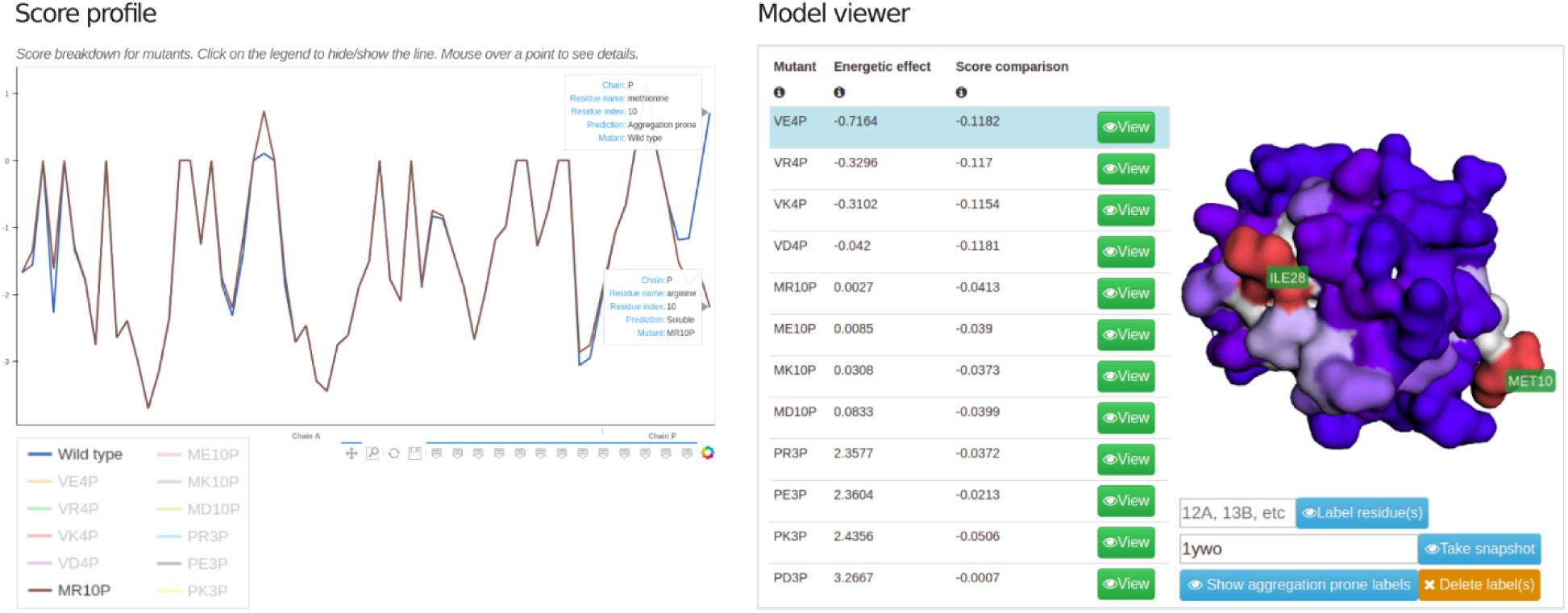
*Automated mutations* tab provides similar functionality to the *Structure* tab but for all the mutants. The additional interactive *Score profile* plot shows the differences between the mutants. Click on the legend to show/hide selected mutant, mouse over the plot to see the details of each residue and use the tool at the bottom to save the plot.

Finally, we provide illustrative examples of using automated mutation feature in Case Studies sections: 4.1, 4.2 and 4.3.

### 3.3. Dynamic mode run

To gain insights into the effect of protein flexibility on the aggregation propensity, dynamic mode can be used. As mentioned earlier CABS needs to be installed for this option to work. For the command line add a *-d* to the basic run example provided in section 3.1 (for the command syntax see Table 3):

*$ aggrescan -v 4 -w a3d_dynamic -i 2gb1 -d*

Please note, dynamic and automatic protein enhancements are incompatible. For the app tick the *Dynamic mode* box.

A new tab will appear in the app’s results page offering a very similar functionality to the automated mutations output. A3D allows a detailed analysis of each flexible model can be analyzed and compared instead. A plot of the RMSF (the measure of protein structure fluctuations during the CABS-flex simulation) will also appear at the bottom of the *Dynamic* tab and additional files will be generated in the working directory:

- *averages* – Json-like formatted file with Aggrescan3D score for each of the CABS-flex generated models;
- *CABSflex_rmsf.csv* - tab separated file with two columns: residue ID (chain letter + residue index, for example A13) and RMSF score calculated by CABS-flex;
- *CABSflex_rmsf.png* - a plot of the RMSF;
- *models.tar.gz* or *model_x.pdb* - all the models generated by CABS-flex (if the GUI i used, the tar.gz archive is extracted);
- *stats.tar.gz* or *model_x.csv* - Aggrescan3D analysis for each of CABS-flex models. The file is formatted similarly to *A3D.csv*. If the GUI is used, the .tar.gz archive is extracted.

The example of using dynamic mode is provided in section 4.3.

### 3.4. Managing projects

All the projects ran with the A3D app can be found in the *Manage project* tab, which allows the users to delete or re-run projects (see Figure 5). There also is an option to add projects that were ran manually from the command line to the Project Manager and enjoy the benefits of the visualization. In order to do so, click on the ‘Add a project*’* button and select the *config.ini* file that was generated with the command prompt.

**Figure 5.**
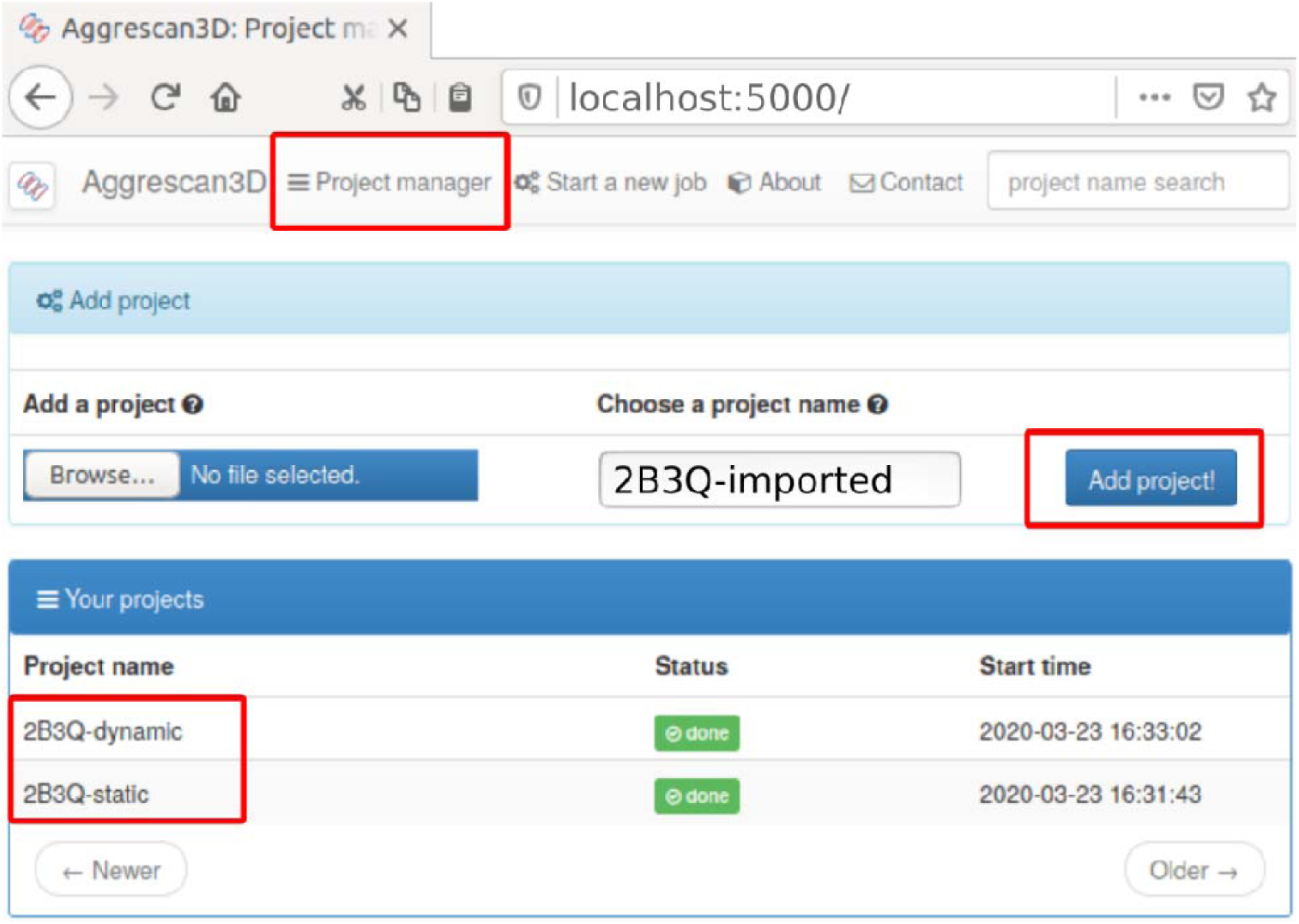
A3D project manager. It contains a list of projects, which results can be accessed by clicking on a project name. If a project was run manually it can still be added to the manager interface.

## 4 Case Studies

Aberrant self-assembly into non-functional aggregates is an intricate molecular process with toxic consequences associated with the onset of different human pathologies^17^. Protein aggregation is also a major and expensive drawback in the manufacturing and storage of protein-based therapies, draining production pipelines and restricting the development of otherwise promising biotherapies^18,19^. Therefore, a large effort has been made to develop efficient strategies for rational design of soluble and stable biotherapeutics, particularly antibodies^20^, and early-stage detection of aggregation-prone regions (APRs). In the case studies below, we discuss the use of the structure-based A3D automated mutation protocol to design mutant’s variants with improved solubility and/or stability without compromising their activity. The case studies include the green fluorescent protein (GFP) (see section 4.1); the heavy chain variable domain (VH) of the human DP47 antibody germline (see section 4.2); and the Fab domain of the monoclonal antibody bevacizumab (Avastin®, Genentech) (see section 4.3). As demonstrated, comparison of theoretically calculated scores with experimental measurements for selected mutants shows that A3D can be effective and low-cost tool supporting rational design of protein-based drugs free from noxious aggregation processes.

### 4.1. Rational design of soluble variants of the green fluorescent protein

For the analysis of the GFP, we used a crystal structure of the folding-optimized GFP (fr-GFP, PDB: 2B3Q^22^). The A3D predictions were obtained using A3D *static* mode with automated mutation (*-am* option) for 10 the most aggregation-prone residues identified by the tool:

*$ aggrescan -i 2B3Q.pdb -w GFP_static -v 4 -f ∼/PATH/FoldX/foldx4 -D 10 -am ‘10 4’*

The A3D analysis for fr-GFP (see left column of Figure 6) indicated several residues with significant aggregation propensity, reported by positive A3D scores, incorporated in a single aggregation-prone patch at the protein surface (V11, G10, Y39, L221) and lesser APR of an isolated valine (see protein surface in the Figure 6). Including mentioned, the 10 most highly scored residues were virtually substituted to charged amino acids. The energetic and solubilizing effects of these mutations are summarized in Table 5. Out of all possible variations (saved in *Mutations_summary.csv* file), the tool automatically selected several the most stabilizing mutations with optimized solubility (highlighted in Table 5).

**Table 5.**
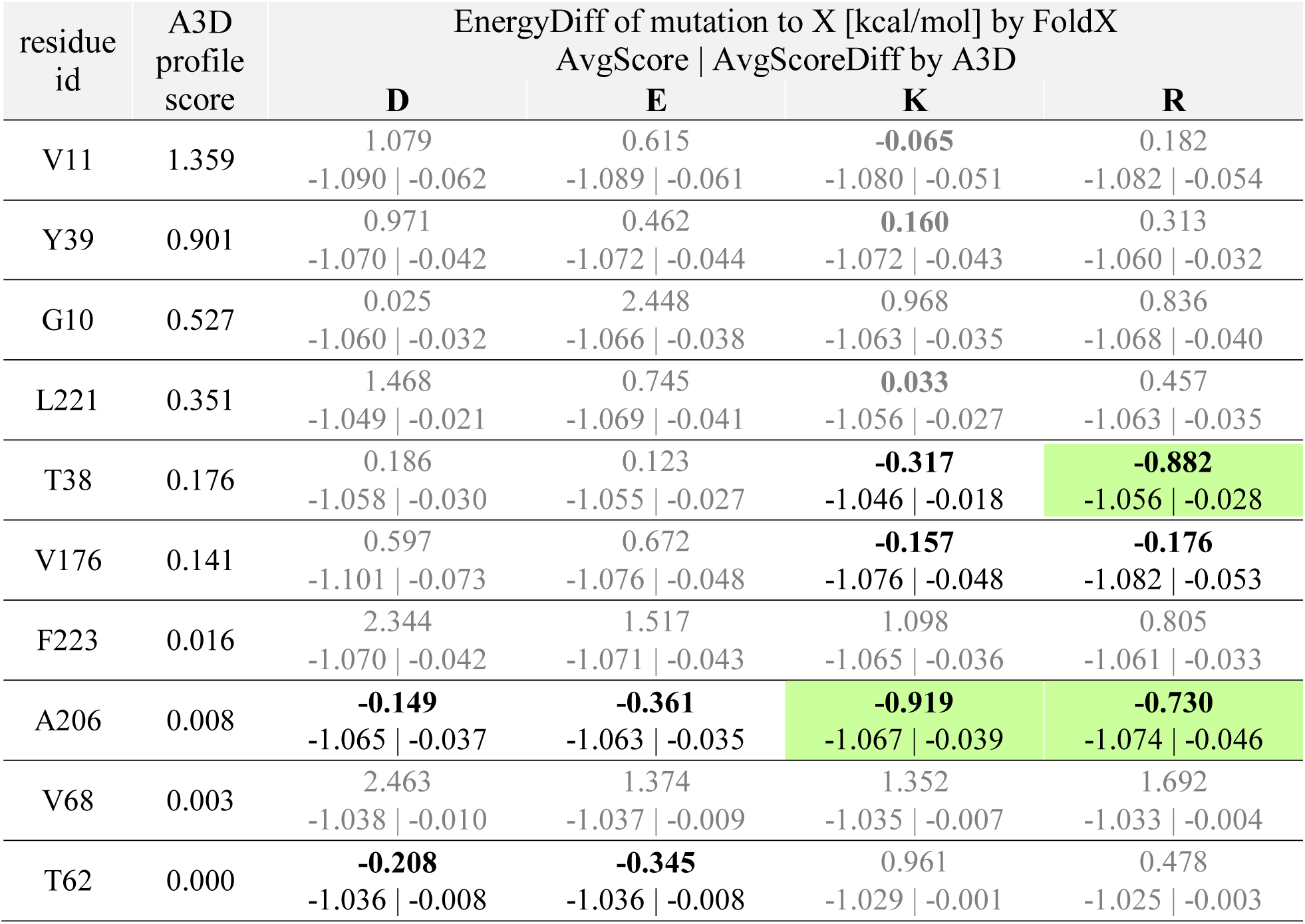
A3D static mode prediction of the automatically generated mutants (-*am* option) in fr-GFP protein. The values in the table correspond to the protein stability (EnergyDiff, stabilizing for ΔG<-0.5, marked in green) and solubility (AvgScore, AvgScoreDiff) parameters. The residues are ordered in descending A3D score and the highest ranked virtual mutants are bolded.

**Figure 6.**
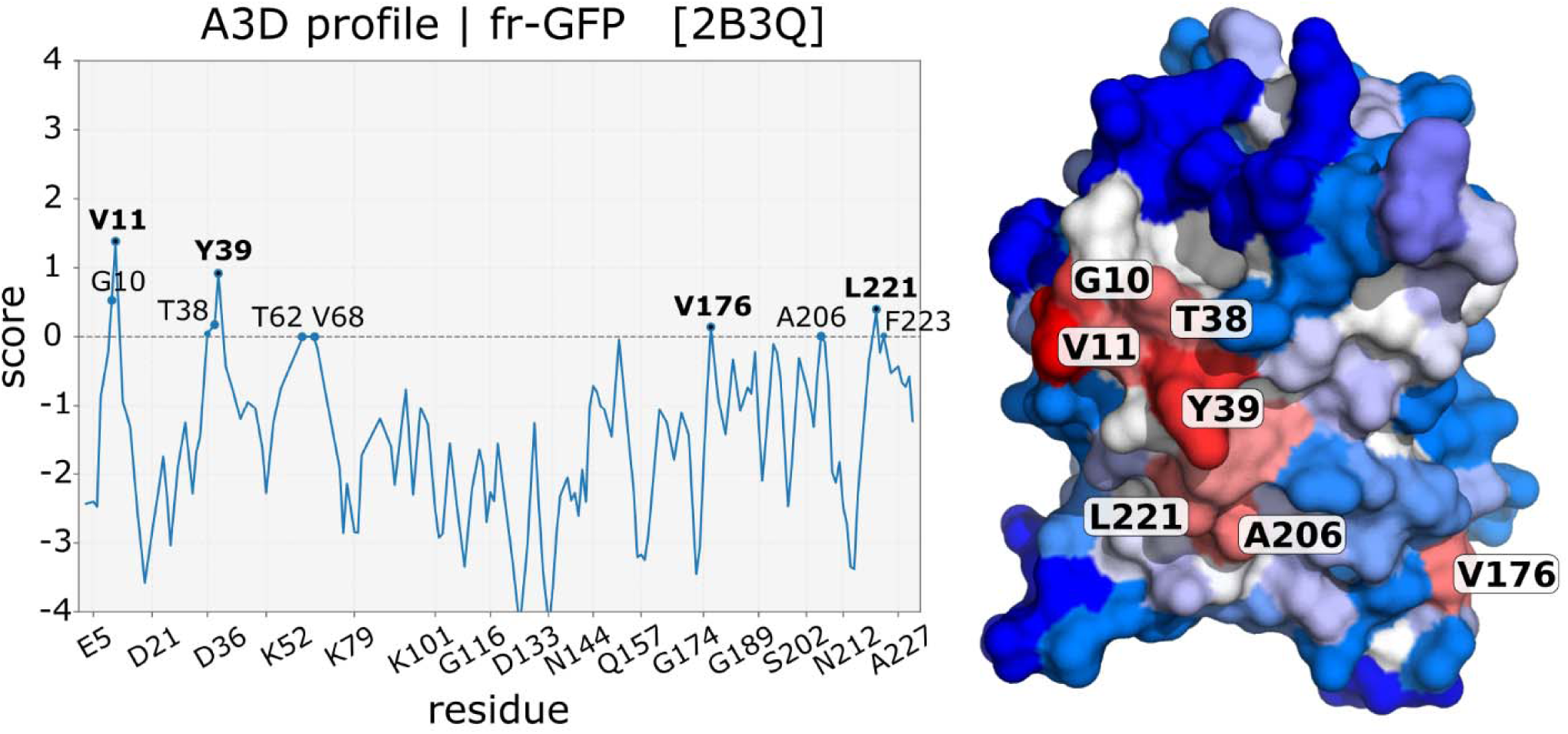
A3D prediction of residue aggregation properties in fr-GFP protein. The left column shows the A3D aggregation profile plot. The right column shows the crystal structure of fr-GFP (PDB: 2B3Q). The protein surface is colored according to A3D score, where blue implies a soluble residues, and predicted aggregation-prone residues are indicated in red shades.

Green fluorescent protein is an example of a protein that contains a modest aggregation-prone region formed by only a few exposed residues. For this reason, when the user requests the A3D screening of virtual mutations on a larger number of the residues, only some of them will have a significant aggregation-prone score, while others may be automatically selected as the most beneficial, in terms of solubility and stability, although being not involved in the structural APRs. That’s the case here. As presented in Table 5, among ten selected residues the highest ranked virtual mutants (black bold font) correspond to residues with the lowest A3D profile score (T62, A206, V176, T38). Moreover, T62 and V68 are not contributing to an exposed APR, being a part of the α-helix that runs through the center of the beta-barrel. F223 and G10 are also important for maintaining stability of the protein spatial structure, which is reflected in the adverse energy effect of all virtual mutations (ΔΔG). Without excluding such residues from the analysis, the user may turn a neutral amino acid into an extra-soluble residue, while maintaining other dangerous aggregation-prone exposed amino acids.

Since the intention in this study is to attenuate the aggregation proneness of the protein, we need to target only the most aggregation-prone residues (V11, Y39, L221) first and then identify the amino acid variation that maximize protein solubility and stability. Note that the same residues would be significant if the user limited the A3D automatic screening to 3-5 top scored residues (default: 2). For these residues A3D scores indicate that mutations to any of the charged amino acid increase protein’s solubility, while their energetic effects may be dramatically different. The introduction of negatively charged residue may have a destabilizing effect (especially Asp mutants), which is a call of special concern for the thermodynamic stability of the protein structure. The positively charged mutants may be significantly more soluble and similarly stable (especially Lys variants) compared to wild type GFP.

Other studies show that the needed effect of increased solubility and stabilization can often only be achieved with the simultaneous introduction of several point mutations^23,24,25^. These predictions can be assessed using A3D with user-indicated mutations (*-m* option). Such exercises, were presented in our previous work^26^, where the effects of cumulative mutations for aggregation-prone residues detected by A3D were analyzed and confirmed experimentally, resulting in the triple mutant GFP/KKK as folded, stable and the most soluble variant (PDB: 6FWW).

### 4.2 Rational design of soluble variants of a single-domain VH antibody

For the analysis of DP47 VH, we used a homology model of DP47 VH that was created by Swiss-Model^27^ based on the 6GHG^28^ template with 98.21% of sequence identity, GMQE (Global Model Quality Estimation) of 0.99 and a QMEAN Z-score of 1.05. The more soluble variants of protein were designed by using A3D default *static* mode with automated mutation (*-am* option) for 10 the most aggregation-prone residues identified by the tool.

*$ aggrescan -i model.pdb -w DP47_static -v 4 -f ∼/PATH/FoldX/foldx4 -D 10 -am ‘10 4’*

The A3D score profile for DP47 VH (see left column of Figure 7) indicated two main groups of aggregation-prone residues: four tyrosines (Y30, Y57, Y98, Y103) and four leucines (L3, L9, L43 and L109). In addition to these, G8, V91 and S97 also scored significantly. The protein surface (see right column of Figure 7), colored as a function of A3D score, shows that all the indicated residues are exposed and contribute to structural aggregation-prone regions. The strongest signal is visible on the APR patch formed by G8, L9, V91 and L109, while L3 is an APR of an isolated amino acid. Finally, ten residues highly ranked by A3D were virtually mutated to charged amino acids. The most stabilizing point substitutions with increased protein solubility are presented in Table 6.

**Table 6.** A3D static mode prediction of the automatically generated mutants (-*am* option) in DP47 VH antibody. The values in the table correspond to the protein stability (EnergyDiff, stabilizing for ΔG<-0.5; no observed) and solubility (AvgScore, AvgScoreDiff) parameters. The residues are ordered in descending A3D score and the highest ranked virtual mutants are bolded. Residues located in CDRs are highlighted in gray.

| residue id | A3D profile score | EnergyDiff of mutation to X [kcal/mol] by FoldX |  |  |  |
| --- | --- | --- | --- | --- | --- |
|  |  | AvgScore \| AvgScoreDiff by A3D |  |  |  |
|  |  | **D** | **E** | **K** | **R** |
| L109 | 1.492 | 1.036<br>-0.557 \| -0.126 | 0.466<br>-0.562 \| -0.130 | **-0.056**<br>-0.556 \| -0.124 | **-0.264**<br>-0.555 \| -0.123 |
| L9 | 1.434 | 0.550<br>-0.535 \| -0.103 | 0.350<br>-0.536 \| -0.104 | **-0.119**<br>-0.532 \| -0.100 | 0.092<sup>11</sup><br>-0.536 \| -0.104 |
| Y98 | 0.998 | 0.118<br>-0.536 \| -0.104 | 0.268<br>-0.537 \| -0.105 | **-0.053**<br>-0.532 \| -0.100 | 0.490<br>-0.537 \| -0.105 |
| Y57 | 0.848 | 1.070<br>-0.529 \| -0.097 | 0.576<br>-0.527 \| -0.095 | **0.038**<br>-0.528 \| -0.096 | **0.026**<br>-0.525 \| -0.093 |
| V91 | 0.838 | 2.419<br>-0.473 \| -0.041 | 1.143<br>-0.469 \| -0.037 | 0.136<br>-0.476 \| -0.044 | 0.235<br>-0.483 \| -0.051 |
| G8 | 0.831 | 2.042<br>-0.482 \| -0.050 | 2.840<br>-0.486 \| -0.054 | 2.0624<br>-0.483 \| -0.051 | 2.257<br>-0.487 \| -0.055 |
| S97 | 0.705 | 1.811<br>-0.446 \| -0.014 | **0.038**<br>-0.453 \| -0.021 | **0.014**<br>-0.457 \| -0.025 | **0.077**<br>-0.470 \| -0.039 |
| W104 | 0.575 | 4.822<br>-0.469 \| -0.038 | 3.769<br>-0.486 \| -0.054 | 2.813<br>-0.479 \| -0.047 | 2.502<br>-0.509 \| -0.077 |
| Y103 | 0.565 | 0.495<br>-0.512 \| -0.080 | 0.201<br>-0.501 \| -0.070 | **-0.066**<br>-0.504 \| -0.072 | 0.162<br>-0.535 \| -0.103 |
| L43 | 0.553 | 2.211<br>-0.529 \| -0.097 | 1.395<br>-0.534 \| -0.102 | 0.853<br>-0.527 \| -0.095 | 1.088<br>-0.541 \| -0.109 |

**Figure 7.**
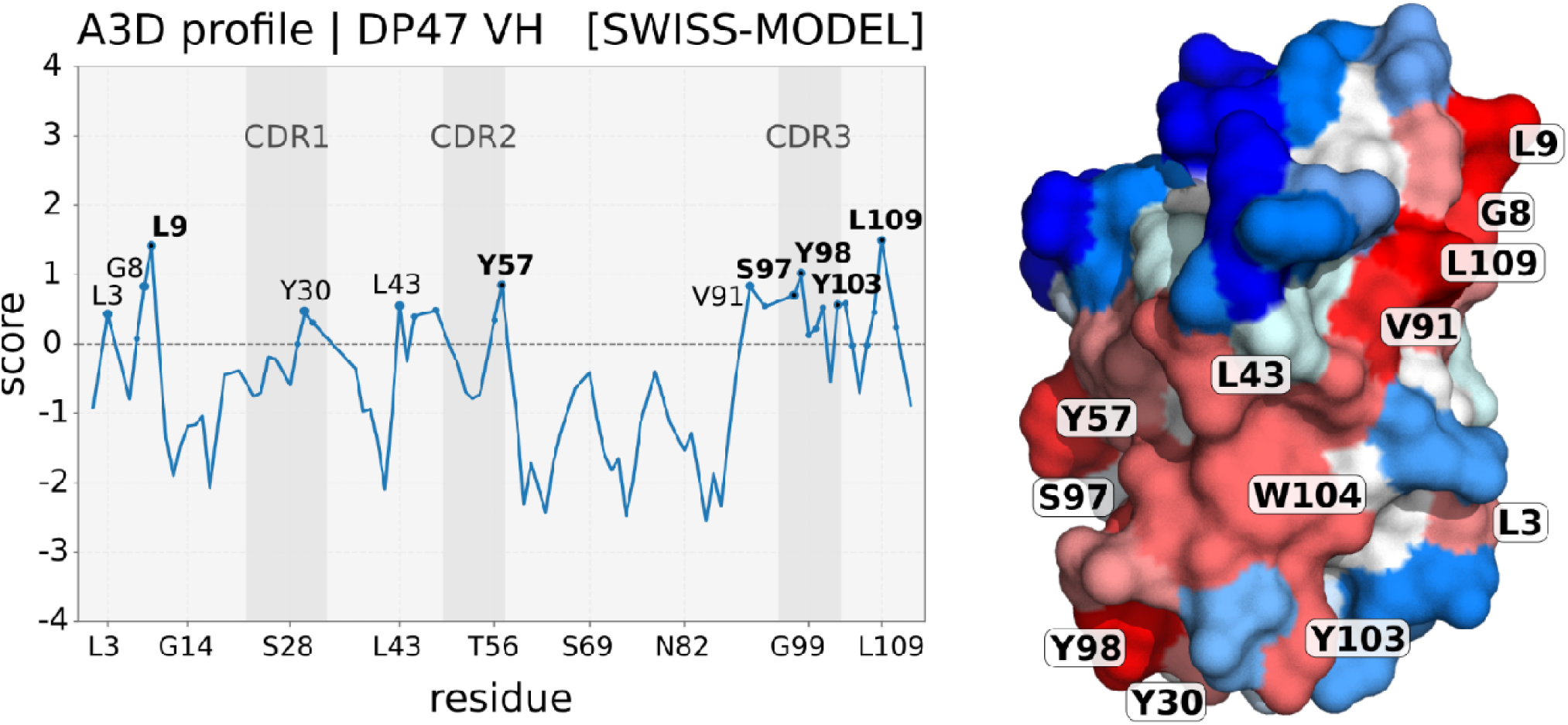
A3D prediction of residue aggregation properties in DP47 VH antibody. The left column show the A3D aggregation profile plot. The right column shows the structure of DP47 model (obtained from SwissModel based on homology template of PDB: 6GHG). The protein surface is colored according to A3D score, where blue implies a soluble residues, and predicted aggregation-prone residues are indicated in red shades.

As presented in Table 6, each of the virtual mutants is predicted to significantly improve the antibody solubility. Despite this generalized protection against aggregation, substitutions to negatively charged amino acids have a destabilizing impact. Alternatively, substitutions to either Lys or Arg are predicted as neutral or stabilizing variants; with the exception of G8, L43 and W104. While any of the substitutions of L109, L9, S97 to a positively charged amino acids might be further considered as faithful candidate to redesign DP47 VH solubility, we would discard the possibility to include neither Y57, Y98 nor Y103 to the study, although A3D suggests some valid modifications. This is because, the aggregation-prone regions containing these tyrosines overlap to a large extent with all three CDRs^26^ (CDR1: A22-S33, CDR2: I49-Y57, CDR3: A96-Y104) of the VH, a particular region of antibodies responsible to bind antigens. Likely, protein regions involved in protein-protein interfaces are enriched in hydrophobic residues, being usually predicted as APRs by A3D and may be automatically included by A3D as solubilizing mutations. Attenuating the hydrophobicity of such important regions might lead to a decreased affinity, poor specificity and ultimately a loss of protein function; all of them, undesired outcomes of reshaping VP47 VH surface. This is a second call of special concern when redesigning protein surfaces for avoiding the creation of variants mutated in functionally involved regions. The aggregation-prone residues being a part of the complementarity-determining regions (CDRs) of antibody (especially functionally relevant) may be manually excluded from the virtual mutation study by using an optional argument to *–am* option provided as a list of residues that will be not considered for mutations:

*$ aggrescan -i model.pdb -w DP47_static -v 4 -f ∼/PATH/FoldX/foldx4 -D 10 -am ‘10 4 30A 57A 98A 103A’*

### 4.3 Rational design of the Fab domain of the therapeutic monoclonal antibody

For the A3D analysis of the Fab domain of the therapeutic monoclonal antibody, we used the crystal structure of PDB: 1BJ1^29^, removing all other chains except H and L. First, we carried out the *static* mode analysis (*stage 1*) with automated mutations to identify the top-ranked aggregation-prone residues of light (L) and heavy (H) chains:

*$ aggrescan -i 1BJ1-HL.pdb -w Fab_static -v 4 -f ∼/PATH/FoldX/foldx4 -D 10 -am ‘10 4 32L 94L 41H 54H 103H 108W’*

Next, in order to study the effect of protein flexibility on aggregation properties, we run the analysis in *dynamic* mode (*stage 2*):

*$ aggrescan -i 1BJ1-HL.pdb -w Fab_dynamic -v 4 -f ∼/PATH/FoldX/foldx4 -D 10 –d*

Accordingly, we obtained a set of models, generated with CABS-flex, reflecting the prevailing structural fluctuations of the Fab domain.

The highest A3D scoring model was selected as the most aggregation-prone conformer in solution and was used as an input to the *static* mode analysis (*stage 3*) with automated mutations to identify alternative mutants to solubilize the protein:

*$ aggrescan -i model-HL.pdb -w Fab_static_alter -v 4 -f ∼/PATH/FoldX/foldx4 -D 10 -am ‘10 4 32L 94L 41H 54H 103H 108W’*

The crystal structure (output of *stage 1*) and the most aggregation-prone model (output of *stage 2*) reflecting conformational fluctuations are compared in Figure 8. The surfaces of both structures are colored according to the A3D score, which facilitates the identification of structural APRs (red shades highlight residues with A3D_score_>0). As expected, the higher *A3D average score* for the CABS-flex generated model (−0.52 compared to -0.71 for 1BJ1) stems from an increase of exposed aggregation prone residues that in normal conditions (static) would be sheltered from solvent. The lower row in Figure 8 shows the RMSF profile (distances between corresponding residues of the superimposed structures) on the left and superimposed backbones of the structures (1BJ1 in red and CABS-flex in blue) on the right.

**Figure 8.**
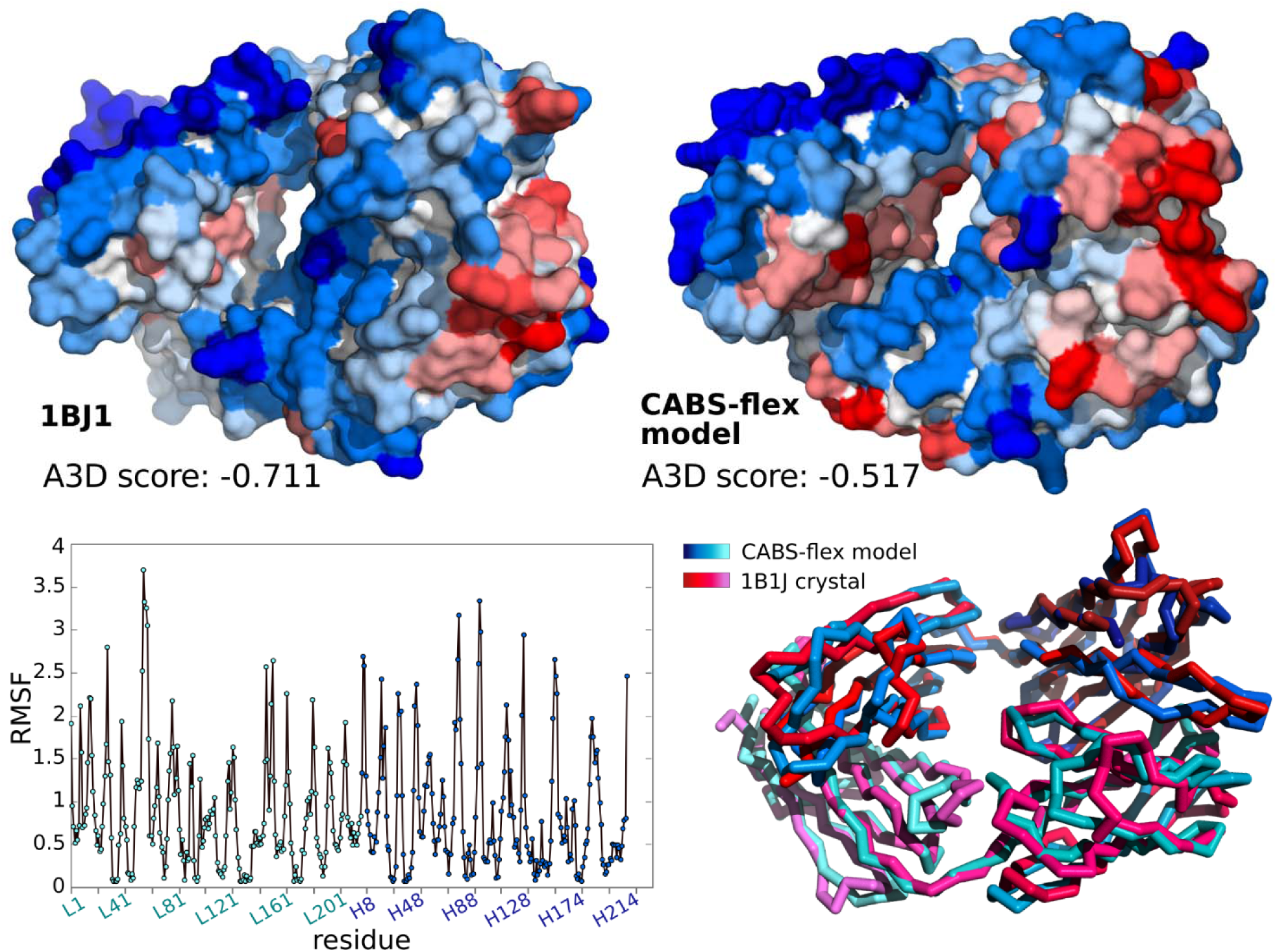
The protein surface colored as a function of A3D score for crystallographic structure of 1BJ1 (top left) and CABS-flex model reflecting conformational fluctuations (top right) of Fab domain. Red shades highlight significant structural APRs, while blue corresponds to soluble residues. The RMSF profile (bottom left) and superimposed backbones (bottom right) of 1BJ1 (red) and CABS-flex model (blue) are provided as a measure of structural flexibility of Fab domain.

The aggregation-prone residues for both the static and the top-ranked CABS-flex model are indicated in the Figure 9, where the upper row shows the protein surface and the lower contains the A3D profiles for H and L chains on the left and right, respectively. Both A3D approaches managed to spot several residues with significant aggregation tendencies. Despite being broadly distributed along the protein sequence, most of these residues colocalize in a specific region of the protein surface and constitute a major structural APR. Based on the available studies^30^ and following an additional visual inspection, we have identified this region as the CDR of the Fab, which comprise the CDR1, CDR2 and CDR3 of the H and L chains. Since the amino acidic composition of the CDR is essential for antibodies to successfully perform their function and bind their targets, we decided to exclude all the residues comprised in this region from the analysis, even though they are scored as highly aggregation prone by A3D (see Y32, Y54, and Y103 from H Chain and Y32, F50, L54 and V94 from L Chain). Of note, residues of CDRs (shaded in grey in aggrescan3D profiles) agglutinate most of the aggregation proneness of the structure, suggesting again that the molecular determinants behind aggregation and inter-molecular interactions are constrained by the same physicochemical properties, such as hydrophobicity.

**Figure 9.**
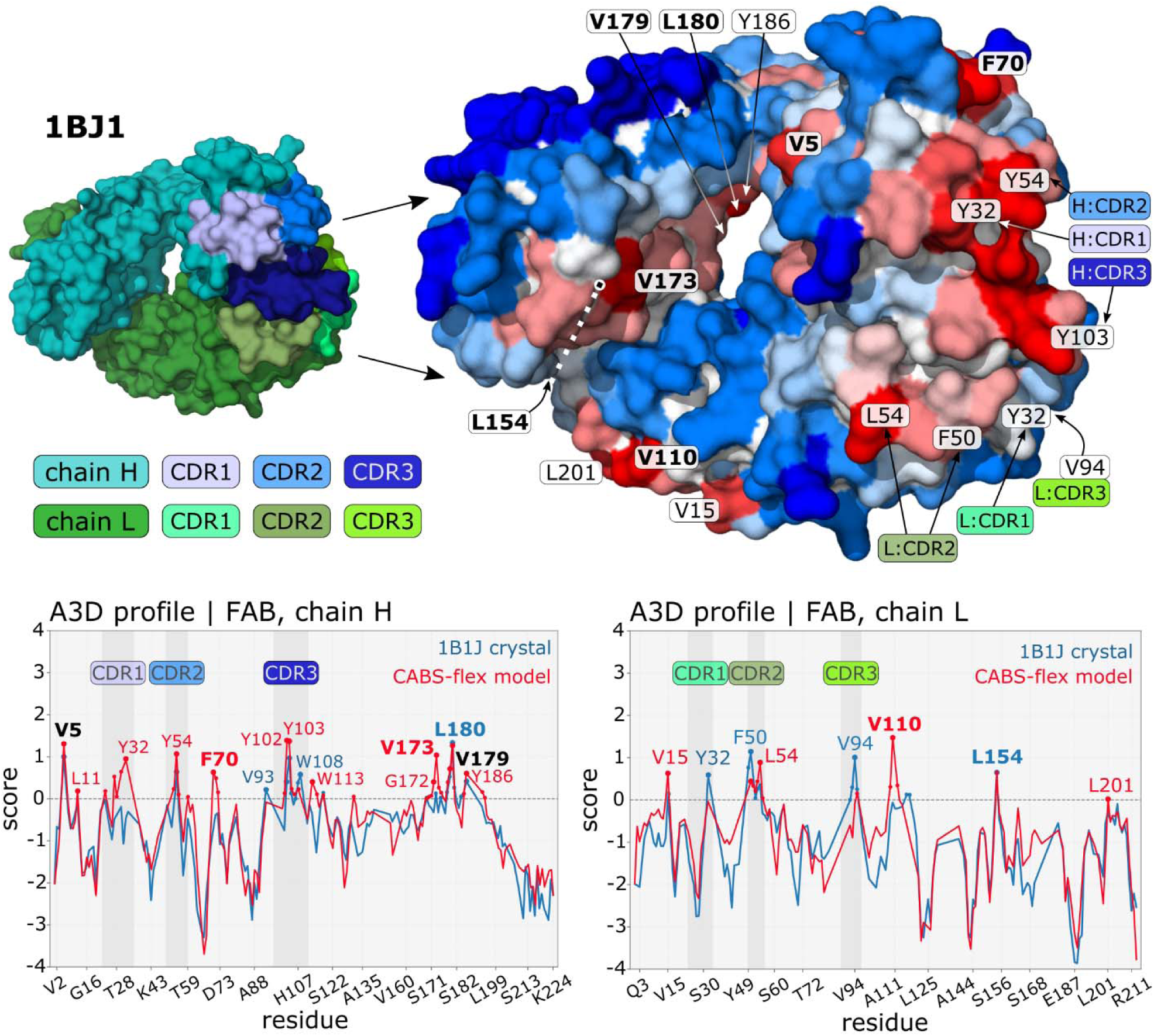
A3D prediction of residue aggregation properties in the Fab domain of monoclonal antibody. The upper row shows: on the left - the crystal structure of Fab domain (PDB: 1BJ1), where H and L chains are indicated in cyan and green, respectively; on the right - the surface of the CABS-flex model, generated in dynamic mode, colored according to A3D score and labeled with highly aggregation-pron residues. The lower row shows the aggregation profile plots for H and L chains on the left and right, respectively. Aggregation-prone residues predicted for 1BJ1 are indicated in blue, while for model reflecting fluctuations in red and common for both structures in black.

Additional residues, not related to CDRs, were also predicted as aggregation prone by A3D, being ideal targets to redesign Fab’s surface. Accordingly, we analyzed solubilizing mutations of both structures with the automated mutations tool of A3D. Table 7 contains the stability (EnergyDiff) and solubility (AvgScore, AvgScoreDiff) values for the A3D suggested mutants. Four of them, V5, V179, L180 from Chain H and L154 from Chain L, were detected by both approaches, which might be explained by a sustained exposure of these amino acids to solvent. In contrast, F70 and V173 from Chain H and V110 from Chain L were exclusively detected in in the most aggregation prone CABS-flex model, probably because their transient exposure to solvent is restricted by structural fluctuations. All computationally created mutants are more soluble than the *wild type* Fab, however substitutions of V5, L180, V110 and L154 residues are predicted to have a greater impact on Fab’s solubility. The effects on structural stability are significantly diverse, although substitutions to positively charged residues are again favored as a general trend, similarly to the previous examples.

**Table 7.**
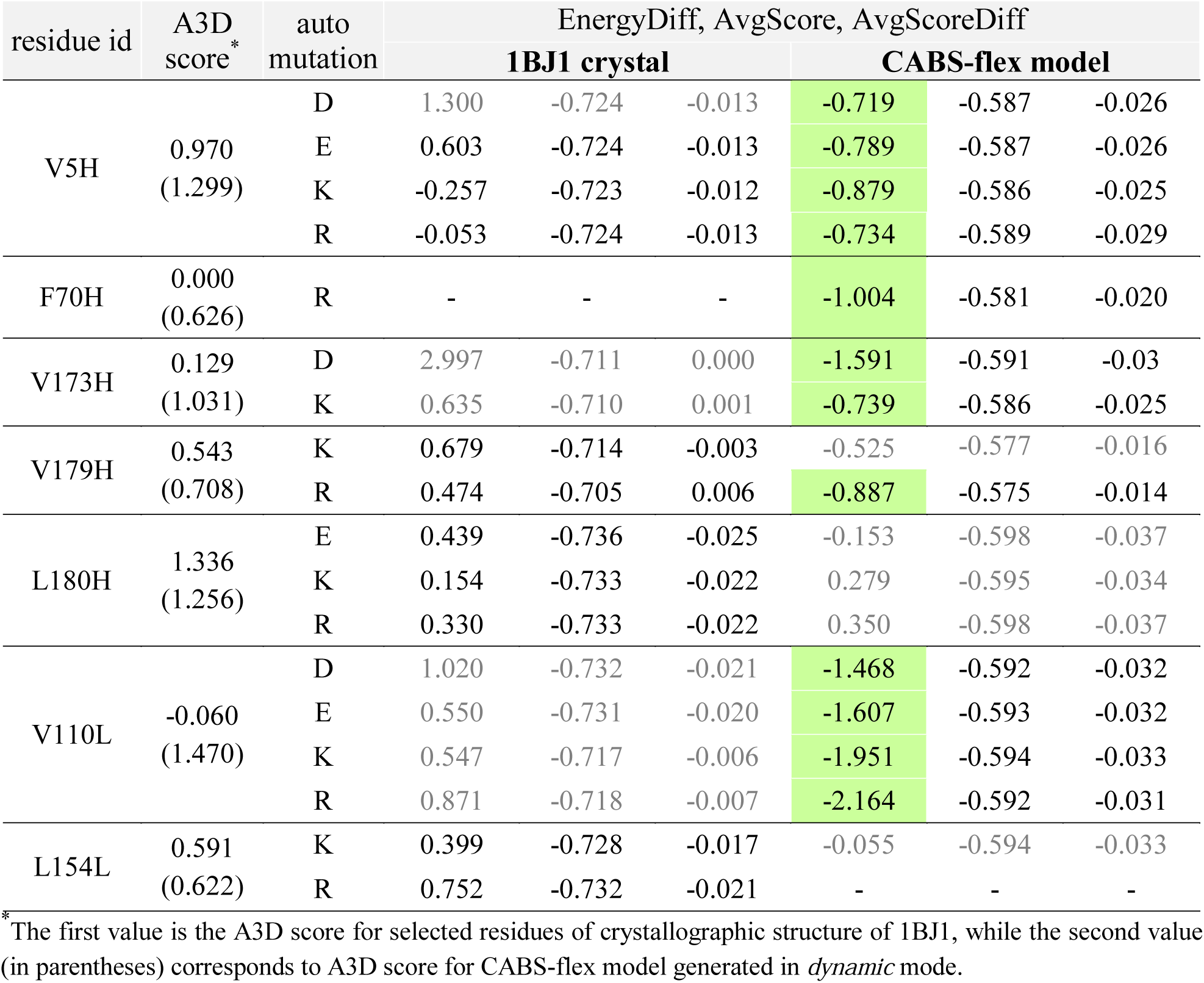
Results for mutations provided by A3D *static* mode with an automated mutations (*-am* option) for the Fab domain. The values in the table correspond to the energy effect (EnergyDiff, stabilizing for ΔG<-0.5, marked in green) of the mutation measured in kcal/mol and calculated by FoldX (embedded in A3D) and solubility effect described by AvgScore predicted by A3D. The grey font corresponds to mutations from outside the highest ranked set.

A recent experimental study performed by Courtois et al.^30^ showed that the same mutations proposed by the A3D automated mutation protocol reduce the aggregation propensity of the fab domain when compared to the wild-type. A3D successfully identified V5, L180, V110 and L154 as the major drivers of aggregation and suggested mutation to K to improve protein’s solubility, while minimizing the impact to protein stability (see Table 8). The implementation of the dynamic mode (*stage 2*) allowed us to spot V110 as an aggregation prone residue, which otherwise would have been unnoticed by the analysis in static mode.

**Table 8.**
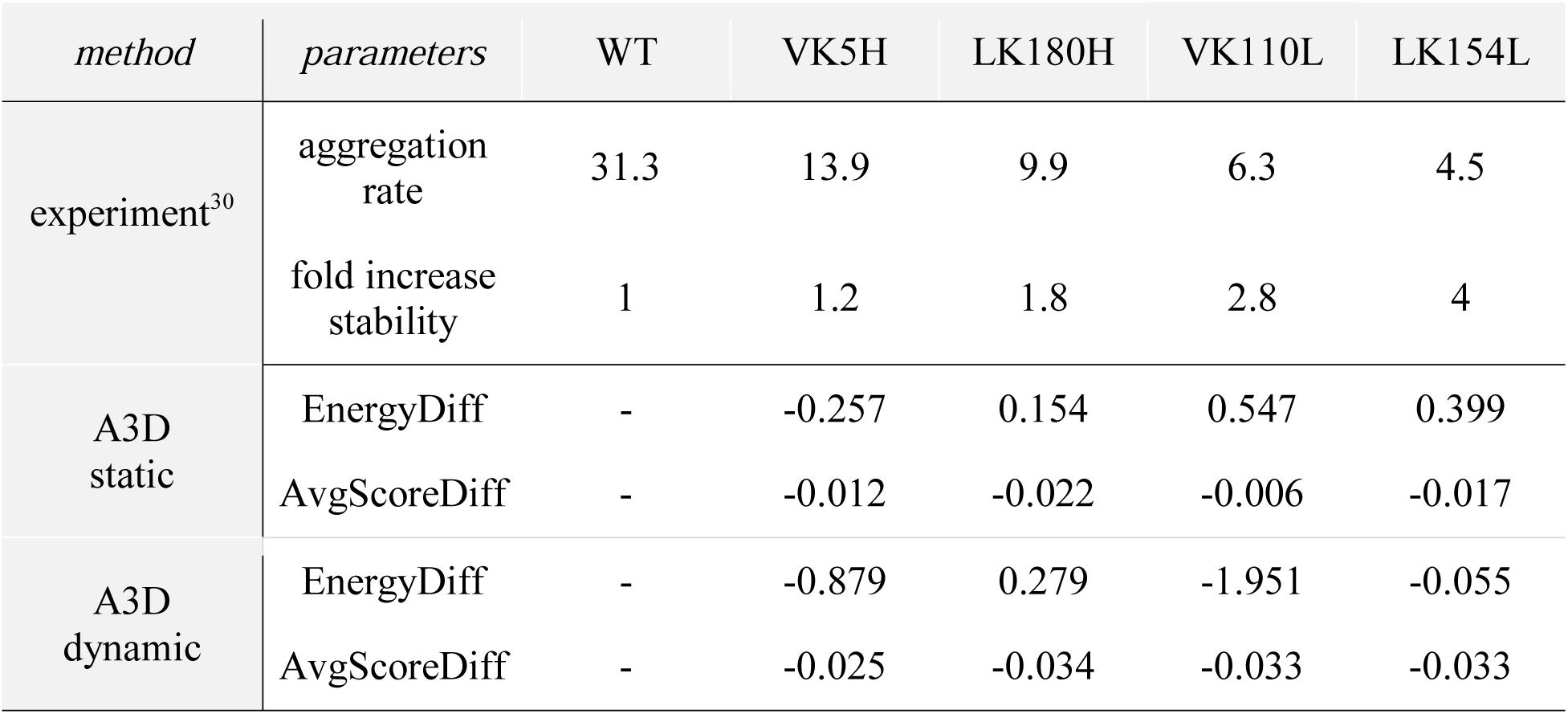
Aggregation properties of the Fab domain variants by experiment and A3D predictions. The table shows experimentally derived parameters^30^ (aggregation rate and stabilization factor “fold increase stability”) and A3D predictions (EnergyDiff and AvgScoreDiff showing changes in stability and solubility, respectively) for the mutants identified in automated mutations A3D prediction run (VK5H, LK180H, VK110L, LK154L).

## 5 Notes

1. Aggrescan3D standalone repository (which contains source code, wiki, installation instructions and issue tracker) is available at https://bitbucket.org/lcbio/aggrescan3d. FoldX suite is available through academic and commercial licenses, details are provided at http://foldxsuite.crg.eu. CABS-flex standalone is distributed under the MIT license, which is free for academic and non-profit users. CABS-flex source code, wiki and installation instructions are available at the repository: https://bitbucket.org/lcbio/cabsflex.
2. Technical hints for A3D options:

3. The mutation code is in the format: <Old Residue type> <New Residue type><Residue number><Chain ID>. An example of a mutation code: ‘VM1A’ means that valine, which is 1st residue of the protein chain A, was mutated into methionine.

| *option* | *useful tips* |
| --- | --- |
| -f, --foldx | Once the option is used with the path */home/user/foldx* it can be omitted in the future runs. |
| -n, --naccess | This option exists mainly to allow for compatibility with the original A3D server which used *naccess*. |
| -m, --mutate | Multiple mutations can be added with a single *-m* line and the *-m* option can be used multiple times. Be careful not to mutate a single |
|  | residue multiple times, this will cause undefined behavior.<br>The mutant is build using FoldX software, alternatively a mutant might be built outside the program and supplied as regular input. |
| -am,<br>--auto_mutation | Automated mutations with *-am* option will choose *N* residues and attempt 4 mutations on each (into glutamic acid, arginine, lysine and aspartic acid). Selecting a large *N* might result in a long simulation.<br>The second argument, *M*, is the number of cores used concurrently to perform the calculations. We advise the number of used cores to be the number of physical cores of your PC or less.<br>An optional argument to *-am* option is a list of residues that will be "fixed" - not considered for mutations.<br>In *dynamic* mode the automated mutation option is not compatible, but *-m* option for user-indicated mutations is available. |
| -c, --config_file | The parenthesis should be omitted in a config file. Detailed instructions on how to create a config file can be found in *configReade.ini* file downloaded with the package.<br>Command line options overwrite those specified in config, so:<br>`$ aggrescan -c config.ini -w /other/dir`<br>will re-run the simulation with a different working directory.<br>Cabs config: Details on how to make those can be found in the CABS program wiki. This gives the user pretty much full control of the CABS simulation. |
| -v, --verbose | **0** - *Only critical* messages will be displayed (usually this means that simulation failed or encountered a serious issue).<br>**1** - *Warning* messages will be displayed as well, these are usually minor issues or situations that might need user's attention.<br>**2** - The default *Info* behavior will display messages about program progress.<br>**3** - *Log files* verbosity level will display messages concerning file management.<br>**4** - *Debug* verbosity level will report as much and accurately as it can about what it is doing.<br>With *-v 4* all the temporary files are left in the 'tmp' folder. These should mostly be useful for debugging purposes. Exceptions will be raised to provide Python traceback, and all temporary files will be kept after the simulation is finished. |

## Acknowledgment

The authors acknowledge support from the National Science Centre of Poland, grant MAESTRO no. 2014/14/A/ST6/00088.

This work was funded by the Spanish Ministry of Economy and Competitiveness BIO2016-78310-R to S.V and by ICREA, ICREA-Academia 2015 to S.V. J.P. was supported by the Spanish Ministry of Science and Innovation via a doctoral grant (FPU14/07161).

